# EPAC1 inhibition protects the heart from doxorubicin-induced toxicity

**DOI:** 10.1101/2021.06.16.448655

**Authors:** Marianne Mazevet, Anissa Belhadef, Maxance Ribeiro, Delphine Dayde, Anna Llach, Marion Laudette, Tiphaine Belleville, Philippe Mateo, Mélanie Gressette, Florence Lefebvre, Ju Chen, Christilla Bachelot-Loza, Catherine Rucker-Martin, Frank Lezoualc’h, Bertrand Crozatier, Jean-Pierre Benitah, Marie-Catherine Vozenin, Rodolphe Fischmeister, Ana-Maria Gomez, Christophe Lemaire, Eric Morel

**Author notes:** These authors contributed equally to the senior authorship of this manuscript (Christophe Lemaire, Eric Morel). deceased (Catherine Rucker-Martin). **Corresponding author: Eric Morel**, UMR-S 1180 – Laboratory of Signaling and Cardiovascular Pathophysiology, Faculté de Pharmacie, Université Paris-Saclay, 17, avenue des Sciences, 91400, Orsay, France.

## Abstract

Anthracyclines, such as doxorubicin (Dox), are widely used chemotherapeutic agents for the treatment of solid tumors and hematologic malignancies. However, they frequently induce cardiotoxicity leading to dilated cardiomyopathy and heart failure. This study sought to investigate the role of the Exchange Protein directly Activated by cAMP (EPAC) in Dox-induced cardiotoxicity and the potential cardioprotective effects of EPAC inhibition. We show that Dox induces DNA damage and cardiomyocyte cell death with apoptotic features. Dox also led to an increase in both cAMP concentration and EPAC1 activity. The pharmacological inhibition of EPAC1 (with CE3F4) but not EPAC2 alleviated the whole Dox-induced pattern of alterations. When administered *in vivo*, Dox-treated WT mice developed a dilated cardiomyopathy which was totally prevented in EPAC1 KO mice. Moreover, EPAC1 inhibition potentiated Dox-induced cell death in several human cancer cell lines. Thus, EPAC1 inhibition appears as a potential therapeutic strategy to limit Dox-induced cardiomyopathy without interfering with its antitumoral activity.

## Introduction

Despite its frequent use and clinical efficiency, the anticancer anthracycline Doxorubicin (Dox) shows strong side effects including cardiotoxicity^1^. Dox cardiotoxicity can be acute with immediate cardiac disorders (arrhythmias) or chronic with remodeling cardiomyopathies (dilated cardiomyopathy (DCM), heart failure (HF)) years after treatment^2^.

Mechanistically, Dox, a DNA Topoisomerase II inhibitor, induces DNA double strand break by direct DNA interaction^3^, oxidative stress^4, 5^, and decreased ATP production^6^. This induces apoptosis and necrosis through p53 cascade and ROS production/ATP depletion respectively^7^ and is proposed to be a primary mechanism of Dox-induced cardiomyopathy^8, 9^. Dox also induces mitochondrial biogenesis alteration and energy defects^10–13^. DNA damage and HF are prevented in cardiac-specific Topoisomerase IIβ (TopIIβ) KO mice, leading to the conclusion that TopIIβ is fundamental in Dox-induced cardiotoxicity^14^. However, alternative mechanisms have been proposed to explain Dox-induced cardiotoxicity^15^. Currently, the only approved medication against anthracyclines cardiotoxicity is dexrazoxane^2^. However, the drug is not devoid of adverse effects, such as potential second malignancies incidence and reduced cancer treatment efficacy^16–18^. Thus, there is a need to search for new therapeutics which would provide long-term cardioprotection against Dox-induced cardiotoxicity without compromising its antitumoral efficacy.

The exchange factor EPAC (Exchange Protein directly Activated by cAMP) contributes to the hypertrophic effect of β-adrenergic receptor chronic activation in a cAMP-dependent but PKA-independent manner^19, 20^. Two EPAC genes are present in vertebrates, EPAC1 and EPAC2^21^. EPAC1 is the main isoform expressed in human hearts and its expression is increased in HF^19^. EPAC is a guanine-nucleotide-exchange factor for the small G-protein Rap^22, 23^ and in pathological conditions, promotes cardiac remodeling through pro-hypertrophic signaling pathways, which involve nuclear Ca^2+^/H-Ras/CaMKII/MEF2 and cytosolic Ca^2+^/Rac/H-Ras/Calcineurin/NFAT^20, 22, 24-26^. Although the role of EPAC in cardiomyocyte apoptosis remains controversial^27–29^, several molecular intermediates in the EPAC1 signalosome (e.g. Ca^2+^ homeostasis, RhoA, Rac) have been shown to be involved in Dox-induced cardiotoxicity^30–32^. There is also evidence that EPAC inhibition may be cardioprotective in ischemia/reperfusion injury^33^. Moreover, EPAC was shown to contribute to cardiac hypertrophy and amyloidosis induced by radiotherapy^34^. Finally, EPAC was shown to play a role in various cancers^35, 36^ and tumoral processes^37–39^, suggesting EPAC as a potential therapeutic target for cancer treatments^36, 40^.

All this suggests that EPAC may play a role in Dox-induced cardiotoxicity. Thus, the aim of this study was to explore this hypothesis by evaluating whether inhibition of EPAC1, either pharmacologically (using the specific EPAC1 inhibitor, CE3F4^41^) or genetically (EPAC1 KO mice), is cardioprotective against Dox-induced cardiotoxicity.

## Results

### Doxorubicin induced DNA damage and mitochondrial caspase-dependent apoptosis in cardiomyocytes

To characterize the Dox induced cell death profile, we performed time response curves (12 h to 48h) of Dox exposure (1 μM) in neonatal rat ventricular myocytes (NRVM) by flow cytometry analysis. Representative monoparametric histograms of FDA fluorescence and TMRM fluorescence, and representative biparametric cytograms for cell size selection, in the presence or absence of EPAC1 inhibitor, are presented in **Extended data Fig. 1a, 1b & 1c**, respectively. **Fig. 1a** shows a time-dependent increase of FDA negative cells, indicative of cell death in association with cell size reduction (**Fig. 1b**), loss of ΔΨm indicative of mitochondrial membrane permeabilization (**Fig. 1c**) and decreased DNA content (**Fig. 1e**). By contrast, Dox did not increase propidium iodide (PI) positive cells percentage, indicator of undamaged cardiomyocytes’ plasma membranes, thus excluding necrosis (**Fig. 1d**). In addition, the level of the active (cleaved) forms of the mitochondrial-dependent initiator caspase 9 and the executive caspase 3 were increased after Dox treatment (**Fig. 1f & 1g**). In the same line, the general caspase inhibitor ZVAD-fmk prevented Dox-induced increase in FDA negative cells and small cells (**Fig. 1h & 1i**), suggesting a caspase-dependent mechanism. DNA damage induced by Dox was assessed by measuring the level of phospho-H2AX (H2AX-pS139, a sensitive marker of DNA double strand breaks) in cultured NRVM at 16 hours to explore the underlying mechanisms prior to the Dox-induced cell death. **Fig. 1j** shows that Dox treatment increased the level of H_2_AX-pS_139_ indicating DNA damage. Altogether, these results demonstrate that Dox induced DNA damage and apoptosis but not necrosis via mitochondrial- and caspase-dependent pathways in NRVM without necrosis induction.

**Fig. 1.**
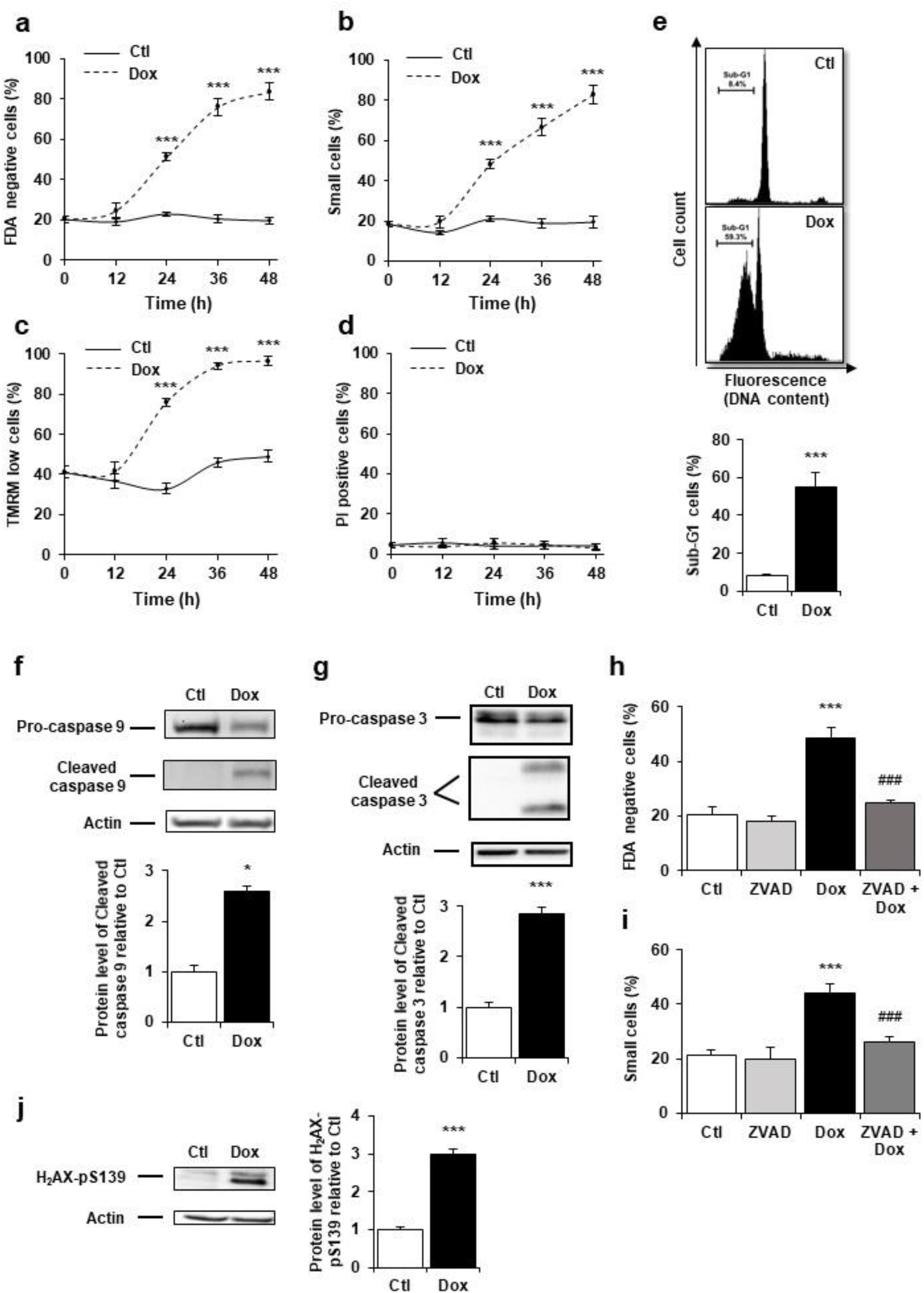
Doxorubicin induces DNA damages and activates mitochondrial pathway of apoptosis in cardiac myocytes. **a to d**, Cell death markers were recorded by flow cytometry in NRVM treated or not with Dox (1 μM) for 12 h, 24 h, 36 h, and 48 h. Results are presented as mean ± S.E.M. (n=12). *** P<0.001 *vs*. control. **a**, Percentages of dead cells (FDA negative cells) measured by FDA assay. **b**, Percentages of small cells obtained by gating the cell population with decrease forward scatter signal. **c**, Percentages of cells with decreased ΔΨm (TMRM low cells) recorded after TMRM staining. **d**, Necrosis was assessed using propidium iodide (PI) and the percentage of cells with permeabilized plasma membrane (PI positive cells) is presented. **e**, NRVM were untreated (Ctl) or treated with Dox (1 μM) for 24 h and the percentage of cells with fragmented DNA (Sub-G1 DNA content) was recorded by flow cytometry and is presented as mean ± S.E.M. (n=4). *** P<0.001 *vs*. control. **f**, **g**, NRVM were treated or not with Dox (1 μM) for 6 h and the levels of cleaved-caspase 9 **f**, or cleaved-caspase 3 **g**, were detected by western blot. Actin was used as a loading control and protein levels relative to Ctl are presented in bar graphs (mean ± S.E.M., n=4 for caspase 9 and n=10 for caspase 3). * P<0.05, *** P<0.001 *vs*. control. **h**, **i**, NRVM were treated or not with Dox (1 μM) +/- the general caspase inhibitor ZVAD-fmk (50 μM) for 24 h. The percentage of dead cells **h**, and small cells **i**, is presented (mean ± S.E.M., n=8). *** P<0.001 *vs*. control, ### P<0.001 *vs*. Dox alone. **j**, NRVM were untreated (Ctl) or treated with Dox (1 μM) for 16 h and the level of the DNA damage marker phosphorylated histone H2AX (H2AX-pS139) was analyzed by western blot. Actin was used as a loading control. Protein levels relative to Ctl are presented in bar graphs (mean ± S.E.M., n=6). *** P<0.001 *vs*. control.

### Dox triggered cAMP-EPAC1 pathway in cardiac cells

To determine whether EPAC1 is involved in Dox-induced cardiotoxicity, we first analyzed EPAC1 protein level and activity after Dox treatment in NRVM. We observed an up-regulation of EPAC1 protein (**Fig. 2a**) by Dox during the acute phase response (3 h and 6 h) which decreased when apoptosis is triggered (24 h). The activity of EPAC1, recorded over time using a pull-down assay of Rap1, showed an increase upon Dox treatment to similar levels as those observed with 10 μM of the EPAC activator 8-CPT (**Fig. 2b**). This increase in EPAC1 activity upon Dox treatment was confirmed using a CAMYEL BRET-sensor assay (**Fig. 2c**), along with an increase of intracellular cAMP concentration, which directly activates EPAC1 (**Fig. 2d**). These results show that Dox induced an increase in both EPAC1 expression and activity in NRVM.

**Fig. 2.**
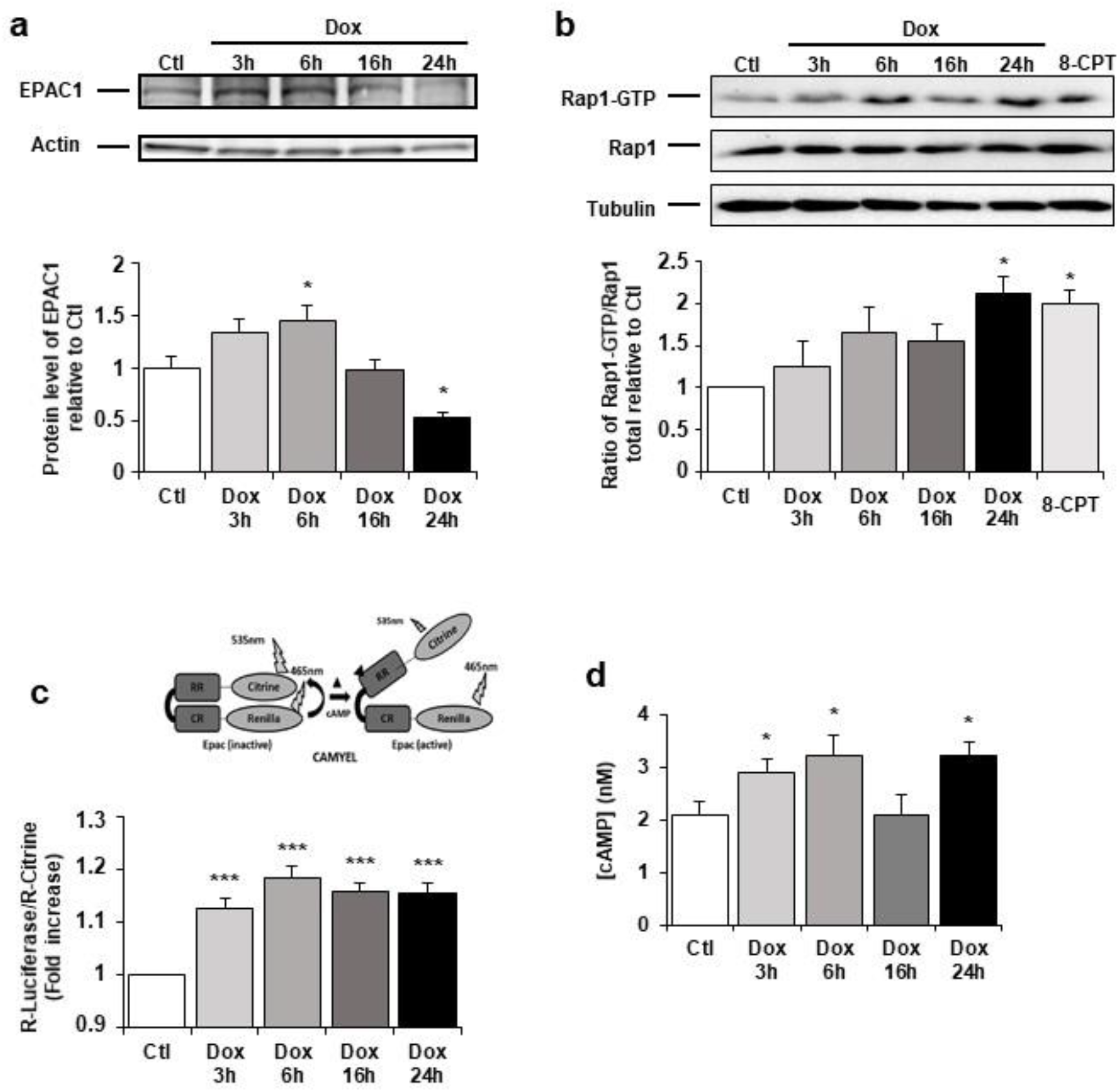
Dox modulates cAMP-EPAC1 pathway in cardiac cells. **a,** NRVM were untreated or treated with 1 μM Dox for 3 h, 6 h, 16 h and 24 h and the level of EPAC1 protein was detected by western blot. Actin was used as a loading control. Protein levels relative to Ctl are presented in bar graphs (mean ± S.E.M., n=7). * P<0.05 *vs*. control. **b**, NRVM were untreated or treated with Dox (1 μM) for 3 h, 6 h, 16 h and 24 h or with the EPAC activator 8-CPT (10 μM) for 3 h. GTP-Activated form of RAP1 was analyzed by pull-down assay. RAP1-GTP/RAP1 total ratios relative to Ctl are expressed in bar graphs as mean ± S.E.M. (n=3). * P<0.05 *vs*. control. **c-d**, NRVM were untreated or treated with Dox (1 μM) for 3 h, 6 h, 16 h and 24 h. **c**, CAMYEL-based EPAC1 BRET sensor was used to measure EPAC1 activation. The BRET ratio was calculated as the ratio of the Renilla luciferase emission signal to that of citrine-cp (means ± S.E.M, n=3). *** P<0.001 *vs*. control. **d**, The concentration of cAMP (nM) was monitored by cAMP dynamic 2 kit (means ± S.E.M., n=4). * P<0.05 *vs*. control.

The role of EPAC in the Dox-induced DNA damages was assessed by using the EPAC1 pharmacological inhibitor, CE3F4^41^. As shown in **Fig. 3a**, CE3F4 (10 μM) decreased H2AX phosphorylation promoted by Dox, suggesting less DNA double strand breaks. Similarly, EPAC1 specific knockdown by shRNA (shEPAC1, **Fig. 3b top panel**) decreased Dox-induced phosphorylation of H2AX (**Fig. 3b lower panel**). Moreover, 8-CPT (10 μM) increased Dox-induced H2AX phosphorylation, while the nonselective EPAC1/EPAC2 inhibitor, ESI-09^42^, but not the selective EPAC2 inhibitor, ESI-05^43^, prevented Dox-induced phosphorylation of H2AX (**Fig. 3c**). These results suggest that the specific inhibition of EPAC1 isoform prevented the formation of Dox-induced DNA double strand breaks.

**Fig. 3.**
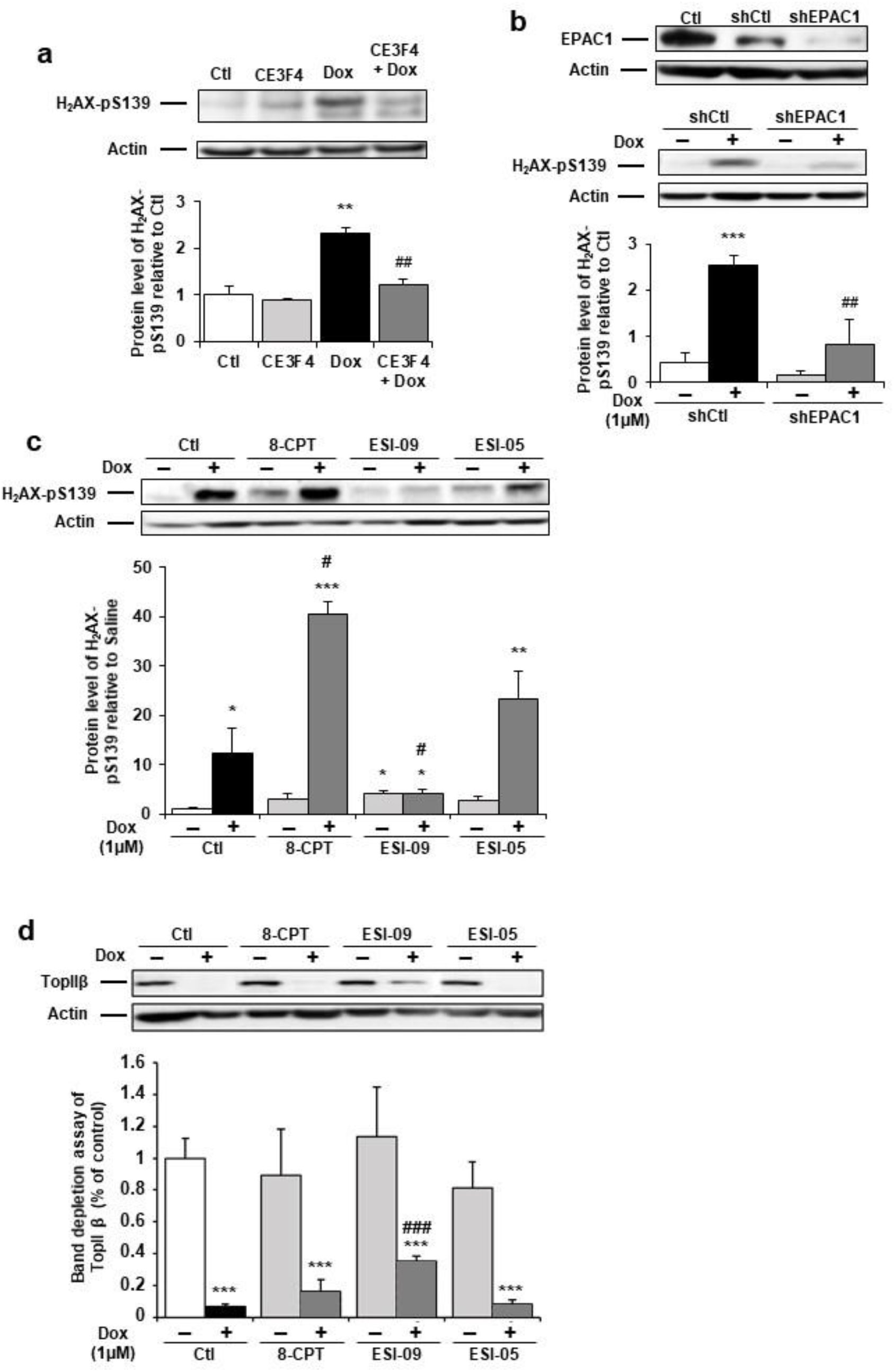
Pharmacological inhibition of EPAC1, but not EPAC2, protects cardiomyocytes from Dox-induced DNA damage. **a**, NRVM were untreated or treated with Dox (1 μM) +/- the specific EPAC1 inhibitor CE3F4 (10 μM) for 16 h and the level of the DNA damage marker H2AX-pS139 was analyzed by western blot. Actin was used as a loading control. Protein levels relative to Ctl are presented in bar graphs (mean ± S.E.M., n=6). ** P<0.01 *vs*. control, ## P<0.01 *vs*. Dox alone. **b**, NRVM were transfected with Control (shCtl) or EPAC1 (shEPAC1) shRNA for 12 h before treatment with Dox for 16 h. The relative levels of EPAC1 and H2AX-pS139 were measured by immunoblotting with actin as a loading control (mean ± S.E.M., n=3). *** P<0.001 *vs*. shCtl, ## P<0.01 *vs*. shCtl+Dox. **c**, **d**, NRVM were untreated or treated with Dox (1 μM) and either the EPAC activator 8-CPT (10 μM) or the EPAC1 inhibitor ESI-09 (1 μM) or the EPAC2 inhibitor ESI-05 (1 μM) for 24 h. **c**, The level of the DNA damage marker H2AX-pS139 was analyzed by western blot. Actin was used as a loading control. Protein levels relative to Ctl are presented in bar graphs (mean ± S.E.M., n=3). * P<0.05, ** P<0.01, *** P<0.001 *vs*. control, # P<0.05 *vs*. Dox alone. **d**, The level of free TopIIβ was quantified by band depletion assay. Actin was used as a loading control. Protein levels relative to Ctl are presented in bar graphs (mean ± S.E.M., n=3). *** P<0.001 *vs*. control, ### P<0.001 *vs*. Dox alone.

TopIIβ has a central role in anthracyclines cardiotoxicity since Dox by stabilizing the cleavable complex DNA/TopIIβ generates DNA double strand break^44^. We therefore examined the EPAC and TopIIβ relation by performing a band depletion assay to quantify free TopIIβ protein, not involved in the DNA/TopIIβ complex (**Fig. 3d**). Dox decreased the free TopIIβ quantity suggesting cleavable complexes formation, an effect in part prevented by ESI-09, but not ESI-05. Altogether, these results indicate that EPAC1, but not EPAC2, is involved in Dox-induced DNA damage and that pharmacologic inhibition of EPAC1 may be useful to prevent Dox-induced DNA/TopIIβ cleavable complex formation and subsequent DNA strand breaks.

### EPAC1 inhibition reduced Dox-induced mitochondrial apoptotic pathway

As metabolism and mitochondrial disorders are an important pattern of Dox cardiotoxicity downstream of TopIIβ signalling alterations, we next investigated whether EPAC1 inhibition protects NRVM from Dox-induced mitochondrial alterations. As shown in **Fig. 4a**, CE3F4 reduced the Dox-evoked dissipation of ΔΨm. Prolonged opening of the MPTP is one of the mechanisms known to initiate mitochondrial membrane permeabilization leading to ΔΨm loss. MPTP opening was thus analyzed by flow cytometry using calcein-cobalt assay. Dox treatment elicited opening of the MPTP (calcein negative cells), which was decreased by EPAC1 inhibition (**Fig. 4b**). We next evaluated the mitochondrial function under Dox ± CE3F4 treatment by measuring the activity of the complex I and IV by cytochrome C oxidase and ubiquinone oxidoreductase activities measurements. EPAC1 inhibition prevented the Dox-induced complex I and IV activity decrease (**Fig. 4c & 4d**). Altogether, these data show that EPAC1 inhibition by CE3F4 protects NRVM from Dox-induced mitochondrial dysfunction.

**Fig. 4.**
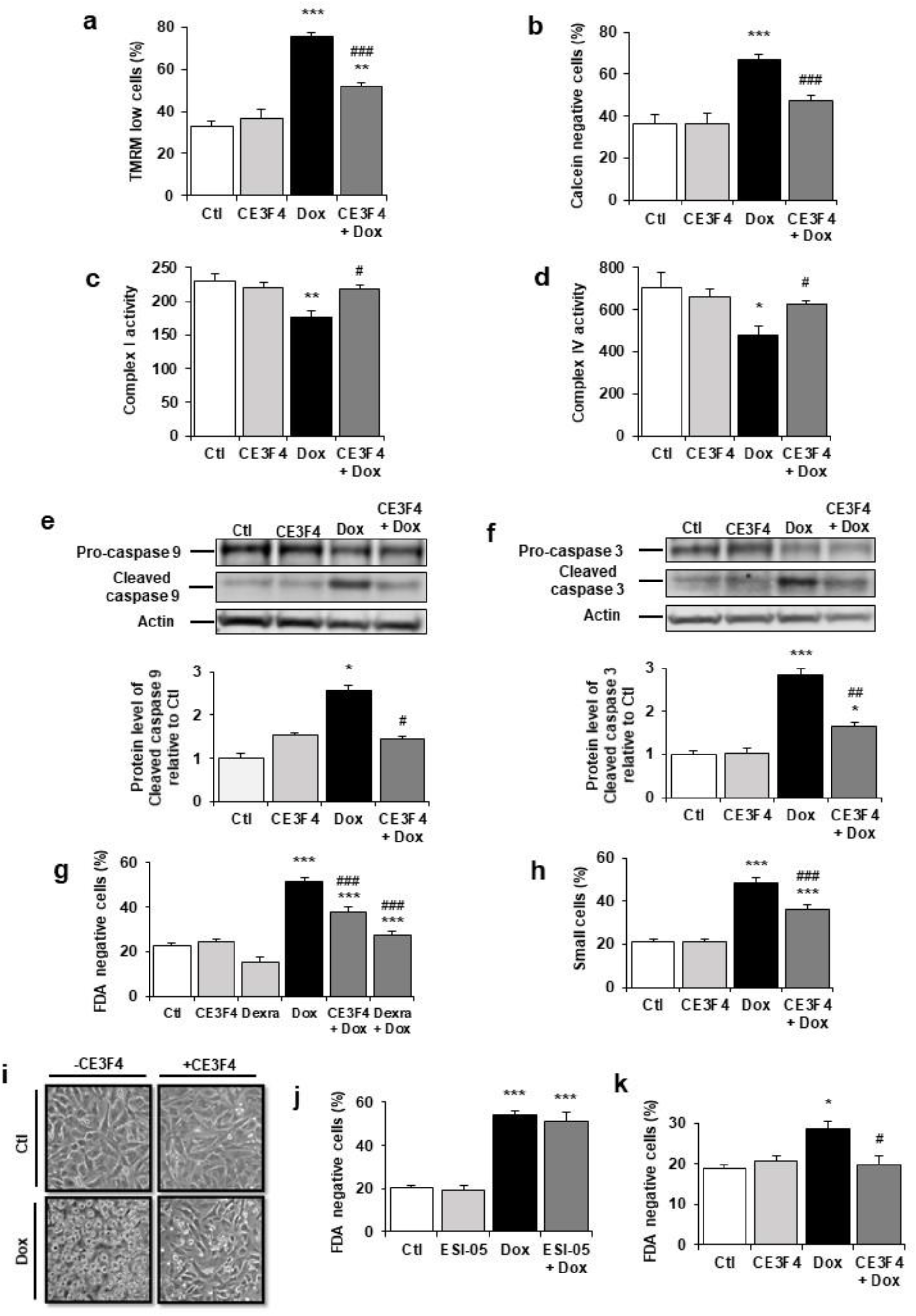
EPAC1 inhibition reduces Dox-induced mitochondrial apoptotic pathway. **a, b**, NRVM were untreated or treated with Dox (1 μM) +/- CE3F4 (10 μM) for 24 h and analyzed by flow cytometry. Results are expressed as means ± S.E.M. (n=4). ** P<0.01, *** P<0.001 *vs*. control, ### P<0.001 *vs*. Dox alone **a**, TMRM staining was used to assess the percentage of cells with decreased ΔΨm (as TMRM low cells). **b**, Calcein-cobalt assay was used to record MPTP opening (as calcein negative cells). **c, d**, NRVM were untreated or treated with Dox (1 μM) and +/- CE3F4 (10 μM) for 16 h. Activity of the mitochondrial respiratory complex I **c**, and complex IV **d**, is presented in the bar graphs as means ± S.E.M. (n=5). * P<0.05, ** P<0.01 *vs*. control, # P<0.05 *vs*. Dox alone. **e, f**, NRVM were untreated or treated with Dox (1 μM) +/- CE3F4 (10 μM) for 6 h and the levels of cleaved-caspase 9 **e**, and cleaved-caspase 3 **f**, were detected by western blot. Actin was used as a loading control and protein levels relative to Ctl are presented in bar graphs (means ± S.E.M., n=4 for caspase 9 and n=10 for caspase 3). * P<0.05, *** P<0.001 *vs*. control, # P<0.05, ## P<0.01 *vs*. Dox alone. **g, h**, NRVM were treated or not with Dox (1 μM) +/- CE3F4 (10 μM) for 24 h and analyzed by flow cytometry. Results in bar graphs are expressed as mean ± S.E.M. (n=12). *** P<0.001 *vs*. control, ### P<0.001 *vs*. Dox alone. **g**, Percentage of dead cells (FDA negative cells). **h**, Percentage of small cells. **i**, Representative micrographs of NRVM untreated or treated with Dox (1 μM) +/- CE3F4 (10 μM) for 24 h. **j**, The percentage of dead cells (FDA negative cells) was assessed in NRVM left untreated or treated with Dox (1 μM) +/- the specific EPAC2 inhibitor ESI-05 (1 μM) and presented as means ± S.E.M. (n=6). *** P<0.001 *vs*. control, ### P<0.001 *vs*. Dox alone. **k**, Freshly isolated ARVM were treated or not with Dox (1 μM) +/- CE3F4 (10 μM) and cell death was determined. The percentage of dead cells (FDA negative cells) is presented in bar graphs (means ± S.E.M., n=4). * P<0.05 *vs*. control, # P<0.05 *vs*. Dox alone.

We also investigated whether EPAC1 is involved in Dox-induced apoptosis. The Dox-induced increase in the level of active caspase 9 (**Fig. 4e**), caspase 3 (**Fig. 4f**), the percentages of FDA negative cells (**Fig. 4g**) and small cells (**Fig. 4h**) were significantly reduced by CE3F4, while ESI-05 did not (**Fig. 4j**). Interestingly, the CE3F4 protection was comparable to that observed with the clinically approved dexrazoxane (**Fig. 4g**). The micrographs shown in **Fig. 4i** depict the morphology of NRVM in the presence or absence of Dox and CE3F4. While many Dox treated-cardiomyocytes aggregated and rounded up, suggestive of dying cells, most CE3F4 supplemented-cells exhibited normal shape. The protective activity of CE3F4 was also tested in a more differentiated model of adult rat ventricular myocytes (ARVM). As in NRVM, CE3F4 protected ARVM from Dox-induced cell death (**Fig. 4k**). These data demonstrate that pharmacological inhibition of EPAC1, but not EPAC2, attenuates Dox-induced cell death both in NRVM and ARVM, suggesting that there is an EPAC isoform specificity in Dox-induced cardiotoxicity.

### Doxorubicin-induced cardiotoxicity was prevented in EPAC1 knock-out mice

Since EPAC1 inhibition showed a strong protection against Dox treatment on various parameters (cell death, mitochondria, DNA damages) in NRVM and ARVM, the obvious question was whether EPAC1 inhibition could also provide cardioprotection *in vivo*. To test this hypothesis, we used mice with global *Epac1* gene deletion (EPAC1 KO). Dox or saline treatment was injected to both WT and EPAC1 KO mice and analyzed 15 weeks after the last injection, when DCM was established^45^. As observed in WT mice (**Extended data Fig. 2a**), a growth delay was observed in Dox treated EPAC1 KO mice in comparison to saline treated mice (**Extended data Fig. 2b**). However, EPAC1 KO mice exhibited a preserved cardiac function after Dox treatment as evidenced by unaltered ejection fraction (EF) and left ventricle end-diastolic volume (LVEDV) (**Fig. 5b-c & 5f-g**) indicating a complete prevention of DCM development. Furthermore, WT heart samples showed that the expression of EPAC1 was increased 6 weeks after Dox treatment and strongly decreased at 15 weeks, when DCM is detected (**Fig. 5d**).

**Fig. 5.**
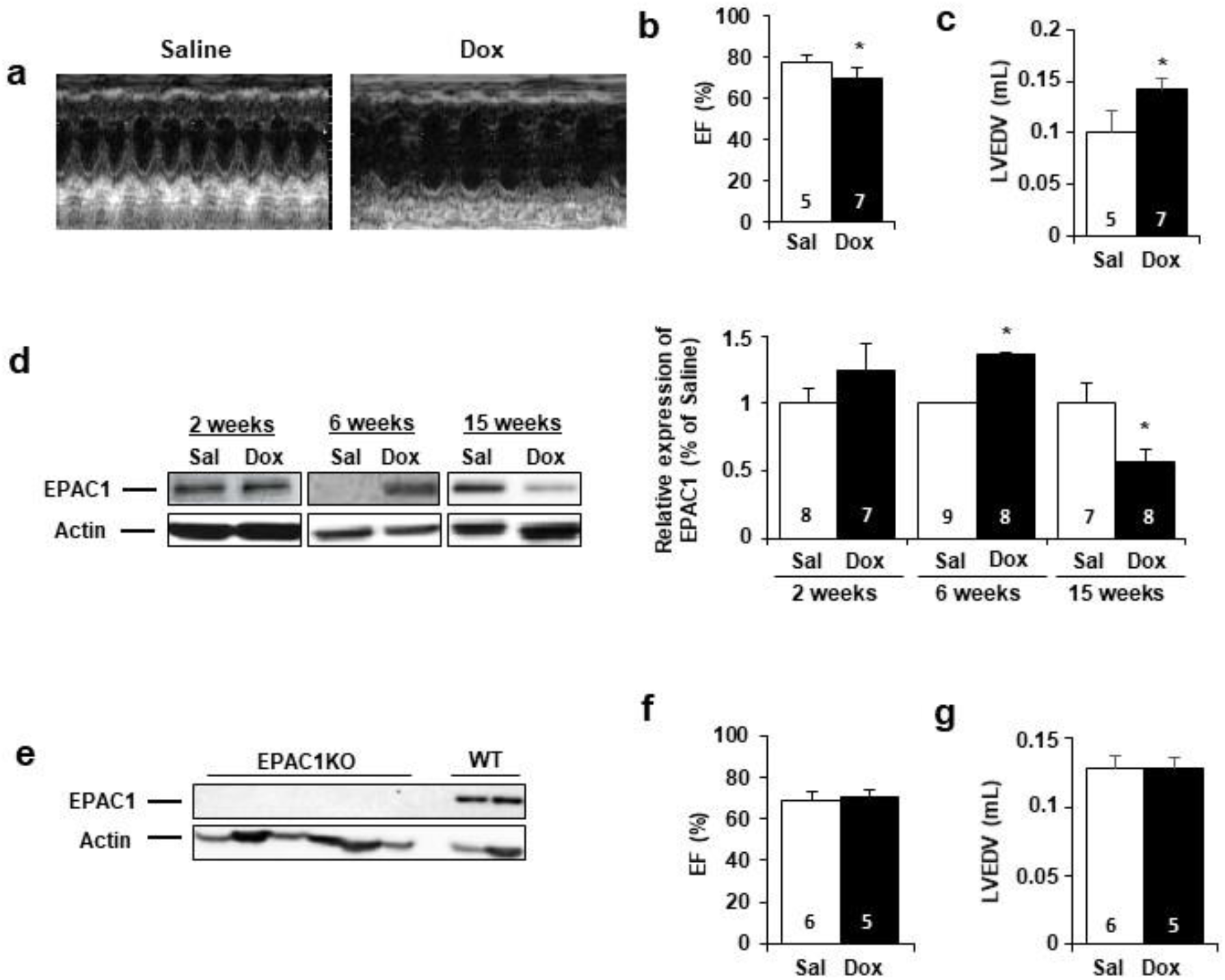
Doxorubicin-induced cardiotoxicity was prevented in EPAC1 KO mice. WT **(a to d)** or EPAC1 KO mice **(e to g)** were injected (i.v.) three times with saline solution (Sal) or Doxorubicin (Dox) at 4 mg/kg for each injection (12 mg/kg cumulative dose). Echocardiographic analysis was performed at 15 weeks after the last injection. **a**, Representative echocardiography images. **b**, Ejection fraction (EF) and **c**, left ventricular end-diastolic volume (LVEDV) are presented as means ± S.E.M. * P<0.05 *vs*. saline. **d**, The protein level of EPAC1 was determined by western blot at 2, 6 and 15 weeks after the last injection. Actin was used as a loading control. Relative protein levels are presented in bar graphs. * P<0.05 *vs*. saline. **e**, Absence of EPAC1 protein was verified in EPAC1 KO mice by western blot. Actin was used as a loading control. **f**, EF and **g**, LVEDV in EPAC1 KO mice are presented as means ± S.E.M.

To verify that the cardioprotection conferred by *Epac1* gene deletion took place at the level of the cardiomyocyte, ventricular myocytes were isolated from WT and EPAC1 KO mice 15 weeks after last i.v. injection and loaded with fluorescence Ca^2+^ dye Fluo-3 AM to record [Ca^2+^]_i_. transients and cell shortening in electrically stimulated cardiomyocytes. **Fig. 6** shows that while cardiomyocytes isolated from Dox-treated WT mice had a reduced unloaded cell contraction (cell shortening, **Fig. 6a**), a reduced [Ca^2+^]_i_. transient amplitude (peak F/F0, **Fig. 6b**), and a decreased sarcoplasmic reticulum Ca^2+^ load (**Fig. 6c**), all these alterations were absent in myocytes isolated from Dox-treated EPAC1 KO mice. **Fig. 6d** also shows that *Epac1* gene deletion prevented the Dox-induced decrease in SERCA2A protein expression. These results demonstrate a complete prevention of Dox-induced toxicity at the level of the cardiomyocyte by genetic deletion of *Epac1*.

**Fig. 6.**
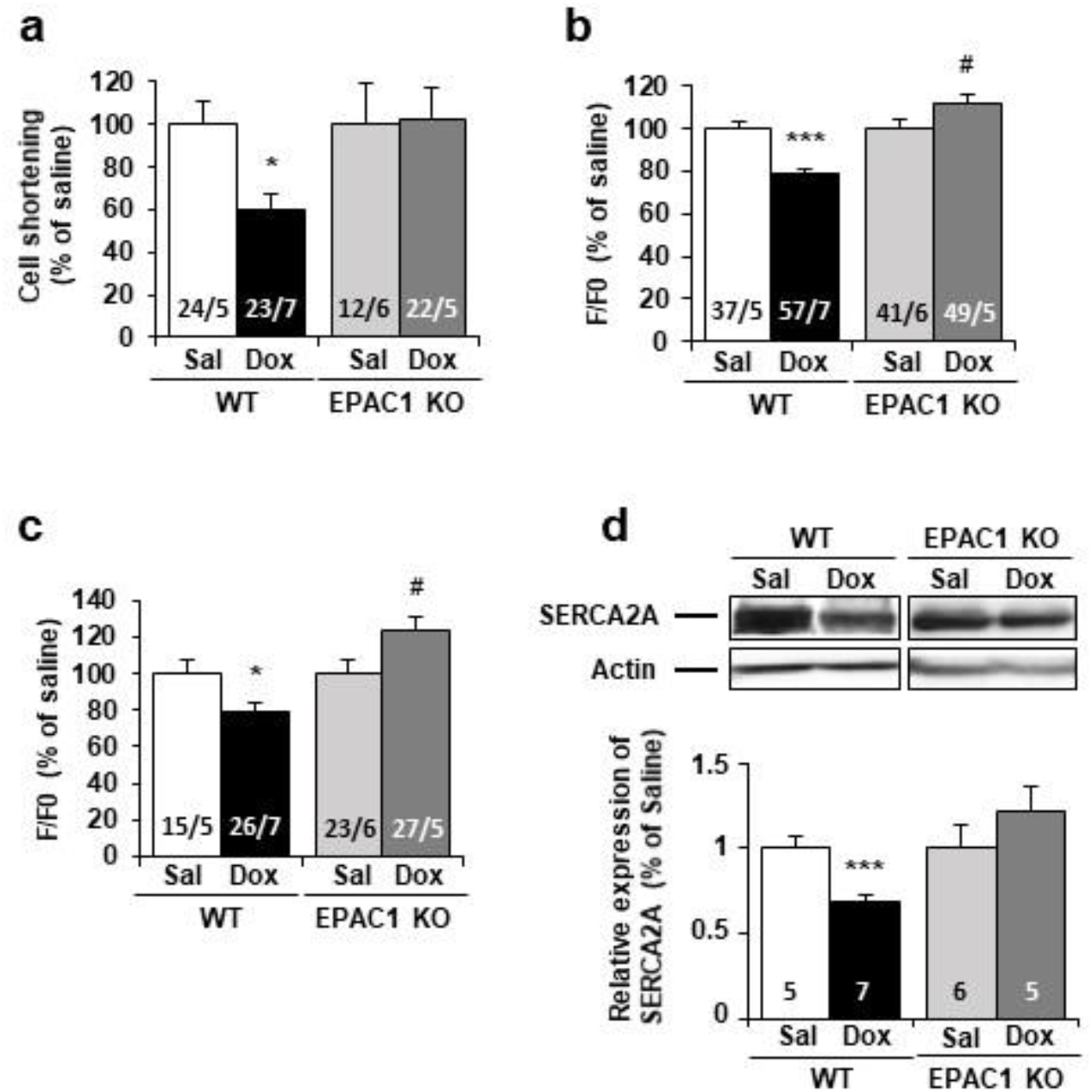
Doxorubicin-induced cardiotoxicity was prevented in EPAC1 KO mice. WT and EPAC1 KO mice were injected (i.v.) three times with saline solution (Sal) or doxorubicin (Dox) at 4 mg/kg for each injection (12 mg/kg cumulative dose). Ventricular cells were isolated from control and treated mice 15 weeks after last i.v. injection and loaded with fluorescence Ca^2+^ dye Fluo-3 AM allowing to measure **a**, cell shortening, **b**, calcium transient amplitude, and **c**, sarcoplasmic reticulum calcium release (upon application of 10 mM caffeine) by confocal microscopy. Normalized values are presented as mean ± S.E.M. * P<0.05, *** P<0.001 *vs*. WT saline, # P<0.05, ## P<0.01 *vs*. EPAC1 KO saline. The number of animals and cells are indicated in the bars of the graphs. **d**, The level of SERCA2A protein was measured by western blot. Actin was used as a loading control. Relative protein levels are presented as mean ± S.E.M. *** P<0.001 *vs*. WT saline. The number of saline or Dox-treated mice are indicated in the bars of the graphs.

### EPAC1 inhibition enhanced the cytotoxic effect of Dox in human cancer cell lines

While the above experiments tend to suggest that EPAC1 may be a potential therapeutic target to limit Dox-induced cardiotoxicity, it is important to verify that EPAC1 inhibition does not compromise the anticancer efficacy of Dox. For that, we tested the effect of CE3F4 on two cancer cell lines, MCF-7 human breast cancer and HeLa human cervical cancer, which are derived from tumors usually treated with Dox. Dox dose response curves ± CE3F4 (10 μM) were performed and cell death (FDA) was measured (after 24 h) using flow cytometry. Dox induced cell death in both cancer cell lines in a dose-dependent manner **(Fig. 7)**. Interestingly, dead cells percentage was significantly increased in both cell lines with CE3F4 (**Fig. 7a & 7b**). Therefore, EPAC1 inhibition not only protects cardiac cells from Dox-induced toxicity but also enhances the anticancer efficacy of this anthracycline.

**Fig. 7.**
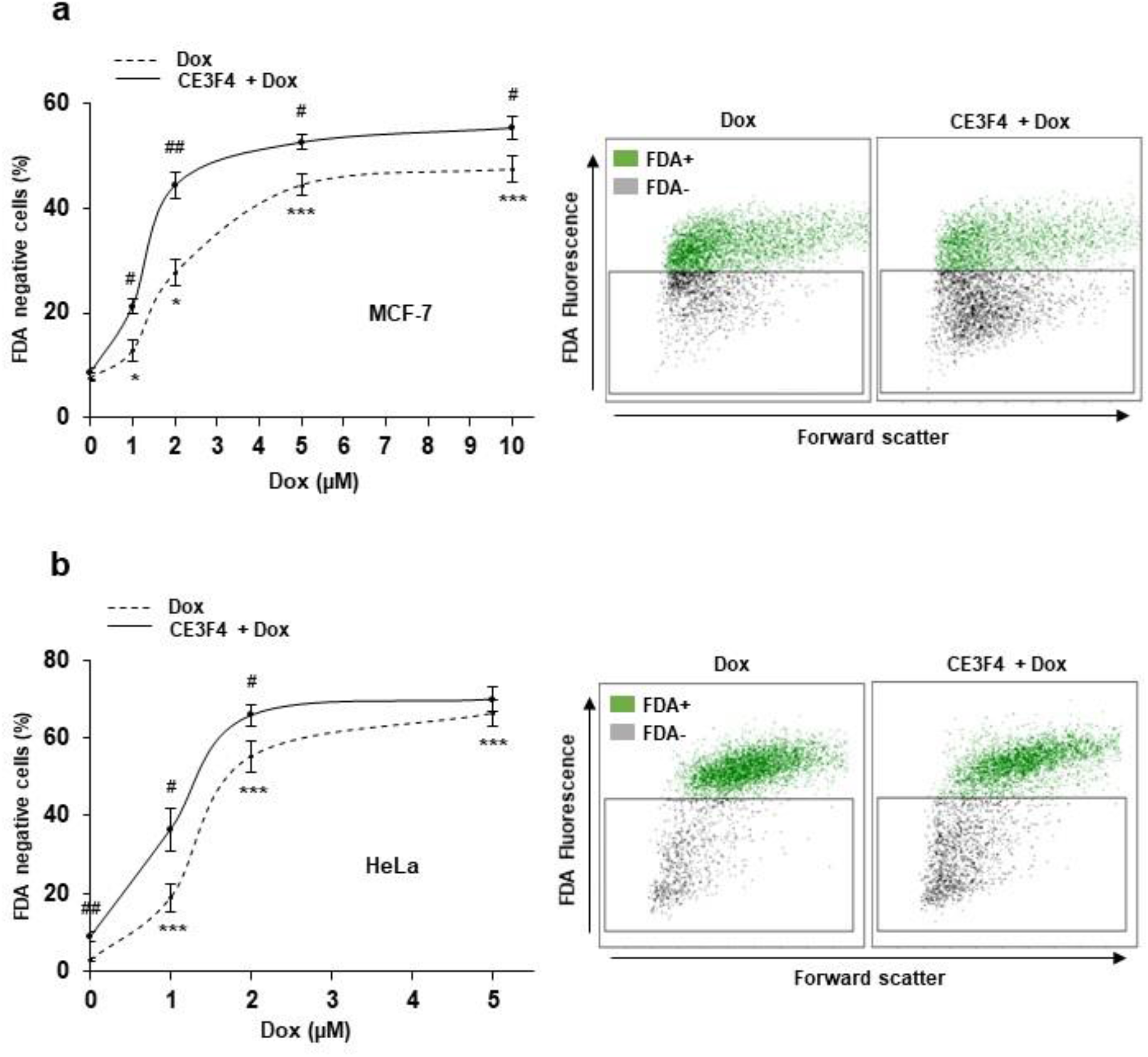
EPAC1 inhibition enhances Dox-induced cytotoxicity in various human cancer cell lines. Human MCF-7 breast cancer cells **a**, and HeLa cervical cancer cells **b**, were untreated or treated with the indicated doses of Dox +/- CE3F4 (10 μM) for 24 h. Cell death was measured by flow cytometry after FDA staining. Results presented in graphs are expressed as the percentage of dead cells (FDA negative cells) (means ± S.E.M., n=5, * P<0.05, *** P<0.001 *vs*. control, # P<0.05, ## P<0.01 *vs*. Dox alone). Representative biparametric cytograms showing FDA fluorescence *vs*. forward scatter (cell size) are presented.

## Discussion

Doxorubicin is a potent chemotherapeutic agent used in clinic to treat a wide range of cancers but this anthracycline is also well-known for its cardiotoxic side-effects. Since decades, Dox-induced toxicity is considered to occur mostly through DNA damage, ROS generation, energetic distress and cell death (apoptosis, necrosis, etc.). However, non-canonical/alternative pathways are nowadays emerging and Dox-induced cardiotoxicity was recently proposed to occur through mechanisms other than those mediating its anticancer activity^14, 46, 47^. Moreover, no widely-accepted therapeutic strategies to minimize Dox-induced cardiac injury have been established. The iron chelator Dexrazoxane is the only FDA and EMEA approved cardioprotector despite limited and still controversial application^18^. Here, we demonstrated that both pharmacological inhibition (CE3F4) or genetic invalidation of EPAC1 prevented Dox-induced cardiotoxicity *in vitro* and *in vivo*. We report an up-regulation of expression and activity of EPAC1 following Dox treatment. Moreover, specific EPAC1 inhibition with CE3F4, but not EPAC2 inhibition with ESI-05, reduced DNA damage, mitochondrial alterations and apoptosis elicited by Dox in cardiomyocytes, indicating that EPAC1, but not EPAC2, inhibition protects the heart against Dox-induced cardiotoxicity. This was confirmed *in vivo* in EPAC1 KO mice. Moreover, EPAC1 inhibition by CE3F4 not only protected cardiac cells, but also increased the toxicity of Dox against human cancer cell lines. Therefore, these results strongly suggest that inhibition of EPAC1 could be a promising strategy to prevent heart damage in anthracyclines-treated cancer patients.

At the molecular level, there is evidence that Dox and EPAC signaling pathways share common features and downstream effectors^30–32^. However, no direct link between these two pathways has been reported so far. Our *in vitro* kinetic analysis revealed that Dox induces an increase of EPAC1 expression in the first hours after treatment (3h-6h), followed by a downregulation (16 h to 24 h). Despite this drop, EPAC activity remained elevated during the 24h of Dox treatment. This was confirmed by the progressive increase of the active form Rap-GTP, the direct EPAC effector, and was most likely due to the observed elevation of cAMP concentration in cardiomyocytes during the first 24 h following Dox treatment. A subsequent decrease of EPAC1 protein level was observed which likely represents a protective cellular response, since we found that chemical or genetic EPAC1 inhibition was cardioprotective.

We show that Dox induces a mitochondrial caspase-dependent apoptosis characterized by DNA damage, Ca^2+^-dependent MPTP opening and mitochondrial membrane permeabilization, caspase activation, nuclear fragmentation and cell size reduction which is consistent with previous report^8^. However, by contrast to other studies^48, 49^, 1 μM Dox did not induce necrosis in NRVM (measured by propidium iodide staining), which was only observed when higher doses of Dox (up to 10 μM) were used (data not shown), indicating that induction of cardiac necrosis by Dox may be dose-dependent.

The role of EPAC1 in cardiomyocyte apoptosis is still unclear owing to controversial reports^27–29, 33^. Indeed, EPAC1 has anti- or pro-apoptotic effect in the heart, depending on the type of cardiomyopathy and its induction mode. For instance, EPAC1 was reported to be cardioprotective and to participate in antioxidant and anti-apoptotic effects of exendin-4, an agonist of the glucagon-like peptide-1 receptor^27^. By contrast, EPAC1 KO mice showed no cardiac damage induced by pressure overload, chronic catecholamine injection or ischemia-reperfusion^28, 33^, suggesting a deleterious effect of EPAC1. This deleterious role of EPAC1 is also found in other tissues as genetic *Epac1* deletion or its inhibition by ESI-09 reduces nerve injury and inflammation, leading to allodynia upon anticancer drug paclitaxel treatment^50^. The protective/detrimental action of EPAC1 is thus not completely understood and might depend on the tissues and the pathophysiological mechanisms of the disease. Here, using pharmacological (CE3F4) and genetic approaches, we found that EPAC1 inhibition is cardioprotective in the context of Dox-induced apoptosis and cardiotoxicity.

Mechanistically, we demonstrated that EPAC1 inhibition prevents MPTP opening induced by Dox. Consistent with our observation, MPTP opening is known to result in mitochondrial membrane permeabilization, pro-apoptotic factors release, activation of caspases and cell dismantling^51, 52^. Recently, it has been demonstrated that EPAC1 KO mice are protected against myocardial ischemia/reperfusion injury^33^. The EPAC1 activation leads to increased mitochondrial Ca^2+^ overload, which in turn promotes MPTP opening, pro-apoptotic factor release, caspase activation and cardiomyocyte death. The demonstration that EPAC1 inhibition prevents Dox-induced MPTP opening and cardiomyocyte death suggests that EPAC1 may play a similar deleterious role in Dox-associated cardiotoxicity and ischemia/reperfusion injury and that MPTP is a major downstream effector in the cardiac cell death signaling cascade regulated by EPAC1.

We found that Dox elicited mitochondrial dysfunction in cardiomyocytes, in agreement with previous report^53^. Indeed, ΔΨm was upregulated together with a strong decrease of the activity of the mitochondrial respiratory complex I and IV in cardiomyocytes exposed to Dox. Importantly, inhibition of EPAC1 by CE3F4 counteracted the mitochondrial alterations induced by Dox. These results could be correlated with a previous study showing that during vascular injury, mitochondrial fission and cell proliferation were suppressed by inhibition of EPAC1 in vascular smooth muscle cells^54^. Therefore, inhibiting EPAC1 can help prevent Dox cardiotoxicity as observed in vascular proliferative diseases.

One of the major mechanisms of Dox-induced cardiomyocyte death and mitochondria reprogramming involves TopIIβ. This gyrase is an enzyme that regulates DNA winding by the formation of DNA/TopII cleavable complexes and generation of DNA double strand breaks^55^. Thus, by its important role in replication and transcription, TopIIβ is of particular interest in cancer therapy^56^. Recently, on the cardiac side, a new paradigm in Dox-induced cardiotoxicity has been put forward, conferring a central role to TopIIβ. Indeed, TopIIβ is required for ROS generation, DNA damage and cell death induction, the three canonical mechanisms involved in Dox side effects^14^. Our results showed that EPAC1 inhibition prevents Dox-induced formation of DNA/TopIIβ cleavable complexes, as evidenced by higher protein level of free TopIIβ, suggesting that EPAC1 inhibition blocks deleterious Dox effects in part through the TopIIβ pathway.

Recent data have reported that EPAC1 could be essential to modulate cancer development and metastasis formation^36^. In addition, inhibition of EPAC1 was proposed as a therapeutic strategy for the treatment of cancers such as melanoma, pancreatic or ovarian cancers^39, 57–59^. For instance, the genetic deletion of EPAC1 or the *in vivo* EPAC1 inhibition by ESI-09 decreases migration and metastasis of pancreatic cancer cells^38^. Of note, in breast cancer cells, which are generally sensitive to Dox therapy, EPAC1 inhibition was shown to inhibit cell migration and induce apoptosis^60^. Here, we found that inhibition of EPAC1 by CE3F4 increased Dox-induced cell death in two human cancer cell lines, including breast cancer cells. Therefore, our data show that EPAC1 inhibition not only protects cardiac cells from Dox-induced toxicity but also enhances the sensitivity of cancer cells to Dox. Similarly, PI3Kγ blockade^46^ or the novel agent named biotin-conjugated ADTM analog (BAA)^47^ were shown to prevent Dox-induced cardiotoxicity and to synergize with its antitumor activity against breast cancer. Therefore, the identification of new promising chemical compound, such as BAA or CE3F4, could be the starting point to the next therapeutic treatments that would protect patients from acute and chronic cardiotoxicity of Dox without altering its cytotoxicity towards cancer cells.

In conclusion, we propose EPAC1 as a new regulator of Dox-induced toxicity in cardiac cells. In response to Dox, the pharmacological inhibition of EPAC1 by CE3F4 recapitulated EPAC1 knock-out phenotype, suggesting the potential therapeutic efficacy of this EPAC1 inhibitor to alleviate Dox-associated side-effects in the heart, while maintaining, or enhancing anticancer effect. Thus, EPAC1 inhibition represents a promising therapeutic strategy both to prevent Dox cardiotoxicity and to enhance its antitumoral activity.

## Methods

### Animal study

All animal experiments were conducted in line with the French/European Union Council Directives for the laboratory animals care 86/609/EEC, (MESRI 18927 authorization).

#### Model

12 weeks old C57BL6 (Wild Type littermate (WT) or *Epac1* Knock-Out (KO)^61^) male mice received *i.v*. (tail vein) NaCl (0.09%) or Dox (4 mg/kg, 3 times at 3 days intervals, cumulative dose of 12 mg/kg) in order to mimic human therapeutic regimen and the induction of a moderated dilated cardiomyopathy without inducing death^62^. Only male have been considered in order to avoid the female hormonal cardioprotection. Cardiac function was measured at 2, 6 and 15 weeks post-injections. Ventricular myocytes were isolated at the same time points for Ca^2+^ handling and Western blot analysis as previously detailed^45^.

#### Echocardiography

Mouse heart physiological parameters were measured by Trans-thoracic echocardiography (Vivid 9, General Electric Healthcare) at 15 MHz, under isoflurane anaesthesia (3%). Left ventricle end-diastolic volume (LVEDV) and ejection fraction (EF) were imaged (two-dimensional mode followed by M-mode). LVEDV was calculated with the Teichholz formula.

### Cell culture

#### Dissociation of Neonatal (NRVM) and Adult (ARVM) Rat Ventricular Myocytes

NRVM were isolated from 1-3 days old Sprague-Dawley rats (40 pups per dissociation) (Janvier, Le Genest-Saint-Isle, France). Ventricles were digested with collagenase A (Roche, Meylan, France) and pancreatin (Sigma Aldrich, St Quentin Fallavier, France), separated by a Percoll gradient and plated (DMEM/medium 199 (4:1) (ThermoFischer Scientific, Les Ulis, France). Langendorff method was used to isolate ARVM as previously described^63^.

#### Human Cancer Cell Lines

MCF-7 and HeLa cell lines were provided by C. Leuter from the Institut Gustave Roussy (Villejuif, France). Cell lines were cultured in DMEM+10% fetal bovine serum and antibiotics.

#### Treatments

Doxorubicin (2 mg/mL) and dexrazoxane were obtained from ACCORD (central pharmacy of Institut Gustave Roussy, Villejuif, France) and Sigma (St Quentin Fallavier, France), respectively. The *in vitro* Dox exposure of isolated cardiomyocytes was of 1-10 μM in concordance with literature recommendation ^6^ and extended to match tumor cell lines exposure requirements. 8-(4-chloro-phenylthio)-2’-O-methyladenosine-3’-5’cyclic monophosphate (8-CPT), ESI-09, and ESI-05 were from Biolog Life Science Institute (Bremen, Germany). ZVAD-fmk was from Bachem (Bubendorf, Switzerland). EPAC1 specific inhibitor (R)-CE3F4 was provided by Dr Ambroise (CEA, Gif-sur-Yvette, France)^64^.

#### Tranfections

Cells were transfected with the GST fusion protein construct or the BRET-based cAMP construct (1 μg) using Lipofectamine 2000 (Invitrogen Life Technologies, Saint Aubin, France) in Opti-MEM medium.

#### Infections

NRVM were incubated for 12 hours with recombinant shRNA encoding shCtl (Welgen^INC^, Worsester, USA, Dr K. Luo) or shEPAC1 (Hôpital universitaire de Nantes, FRANCE, Dr C. Darmon) adenoviruses (MOI 500).

### Cell death profiling

Cell death and apoptotic markers were recorded by flow cytometry (FC500, Beckman Coulter, Villepinte, France). Cell viability was analyzed by FDA (Fluorescein di-acetate) staining (0.2 μg/mL, 10 minutes) in which FDA negative cells were dead cells and those with decreased forward scatter signal counted as small cells entering the apoptotic process. Mitochondrial membrane potential (ΔΨm) was monitored using the cationic dye TMRM (10 nM, 20 min). TMRM low cells were cells with decreased ΔΨm. Necrosis was evaluated by staining cells with 10 μg/mL of Propidium Iodide (PI). PI positive cells were necrotic cells. DNA fragmentation (Sub-G1 DNA content) was estimated by measuring DNA content after overnight permeabilization (70% ethanol) and incubation (4°C, 24h) with PI (50 μg/mL) and RNAse A (250 μg/ml). Mitochondrial Permeability Transition Pore (MPTP) opening was assessed by Calcein-Cobalt assay as previously described^52^.

### Western blot analysis

Proteins were separated by SDS-PAGE (10-15%) and transferred to a PVDF membrane, which were incubated (overnight, 4°C) with antibodies: anti-EPAC1 (1/500), anti-Caspase 3 (1/500), anti-Caspase 9 (1/1000), anti-RAP1 (1/1000), and anti-H2AX-pS139 (1/1000) from Cell Signaling Technology (Saint-Cyr-L’Ecole, France), Topoisomerase II β (1/1000) from Abcam (Paris, France), SERCA2A (1/1000) and anti-Actin (1/50000) from Santa Cruz Biotechnology (Heidelberg, Germany), and anti-Tubulin (1/1000) from Sigma (St Quentin Fallavier, France). Proteins were detected on the iBright FL1000 Imager (ThermoFischer Scientific, Les Ullis, France) with ECL and protein band intensity was calculated using ImageJ software and normalized by Actin.

### Measurement of EPAC1 activity and cAMP concentrations

#### Pull-Down assay of RAP1

To measure EPAC1 activity we performed RAP1 Pull-Down experiments using a GST fusion protein containing the RAP1 binding domain of RAL-GDS as previously described^65^.

#### EPAC-based BRET sensor assay (CAMYEL sensor)

EPAC1 activity was assessed using CAMYEL sensor^66^, a BRET-based cAMP construct, composed of EPAC1 sandwiched between Renilla Luciferase and Citrine. NRVM were lysed, centrifuged (20000g, 10 min) and supernatant was treated (RT, 5 min) with 2 μM Coelenterazin before measure. Emission from Renilla Luciferase and Citrine was measured simultaneously at 465 nm and 535 nm in a plate-reader (Beckman coulter, Villepinte, France). EPAC1 activation decreases BRET signal (465 nm/535 nm ratio).

#### cAMP concentration

Cardiomyocytes cAMP concentration was assessed with cAMP dynamic 2 kit (Cisbio, Saclay, France), according to the manufacturer instructions.

### Calcium handling measurement

Cells were loaded with Fluo-3 AM as previously described^45^. [Ca^2+^]_i_ transients were recorded following electrical stimulation at 2 Hz (two parallel Pt electrodes). Images were obtained with a laser scanning confocal microscope (TCS SP5X, Leica Microsystems, Nanterre, France) equipped with an x40 water-immersion objective in the line scan mode (1.43 ms/line). Fluo-3 AM was excited with a white laser fitted at 500 nm, and emission measured at wavelengths above 510 nm. Image analyses were performed by IDL 8.2 software (Exelis Visual Information Solutions, Inc.) and homemade routines^45^.

### Quantification of Topoisomerase IIβ/DNA complex

The formation of TopIIβ/DNA complex was assessed by Band Depletion Assay, as previously described^67^.

### Mitochondrial respiratory chain complexes activities

NRVM were homogenized in ice-cold HEPES buffer and incubated (4°C, 1h) for extraction.

#### Mitochondrial respiratory complex I (NADH: ubiquinone oxidoreductase) activity

Complex I activity was measured by recording the decrease in absorbance at 340 nm caused by oxidation of NADH to NAD^+^ in phosphate buffer supplemented with Decyl-Ubiquinone. In a second step, Rotenone was added to compare total NADH oxidation in cells and NADH oxidation due to complex I activity.

#### Mitochondrial respiratory complex IV (cytochrome C oxidase) activity

Cytochrome C oxidase activity was assayed by the decrease in absorbance at 550 nm caused by oxidation of ferrocytochrome C to ferricytochrome C by cytochrome C oxidase in phosphate buffer.

### Statistics

Results are expressed as mean ± SEM. Differences between 2 groups have been analyzed by non-parametric Mann-Whitney test. The comparison between more than 2 groups was analyzed by Kruskal Wallis test followed by *post hoc* test with Bonferroni correction. Differences were considered significant at *p<0.05, **p<0.01, and ***p<0.001 vs Ctl and # vs Dox alone.

## Acknowledgments

We are grateful to Dr E. Hirsch and Dr A. Ghigo (University of Torino, Italy) and Dr D. Hilfiker-Kleiner (Marburg University Medical School, Germany) for providing cancer cell lines, and to V. Domergue and the IPSIT platform for animal housing.

## Extended data

**Extended data Fig. 1.**
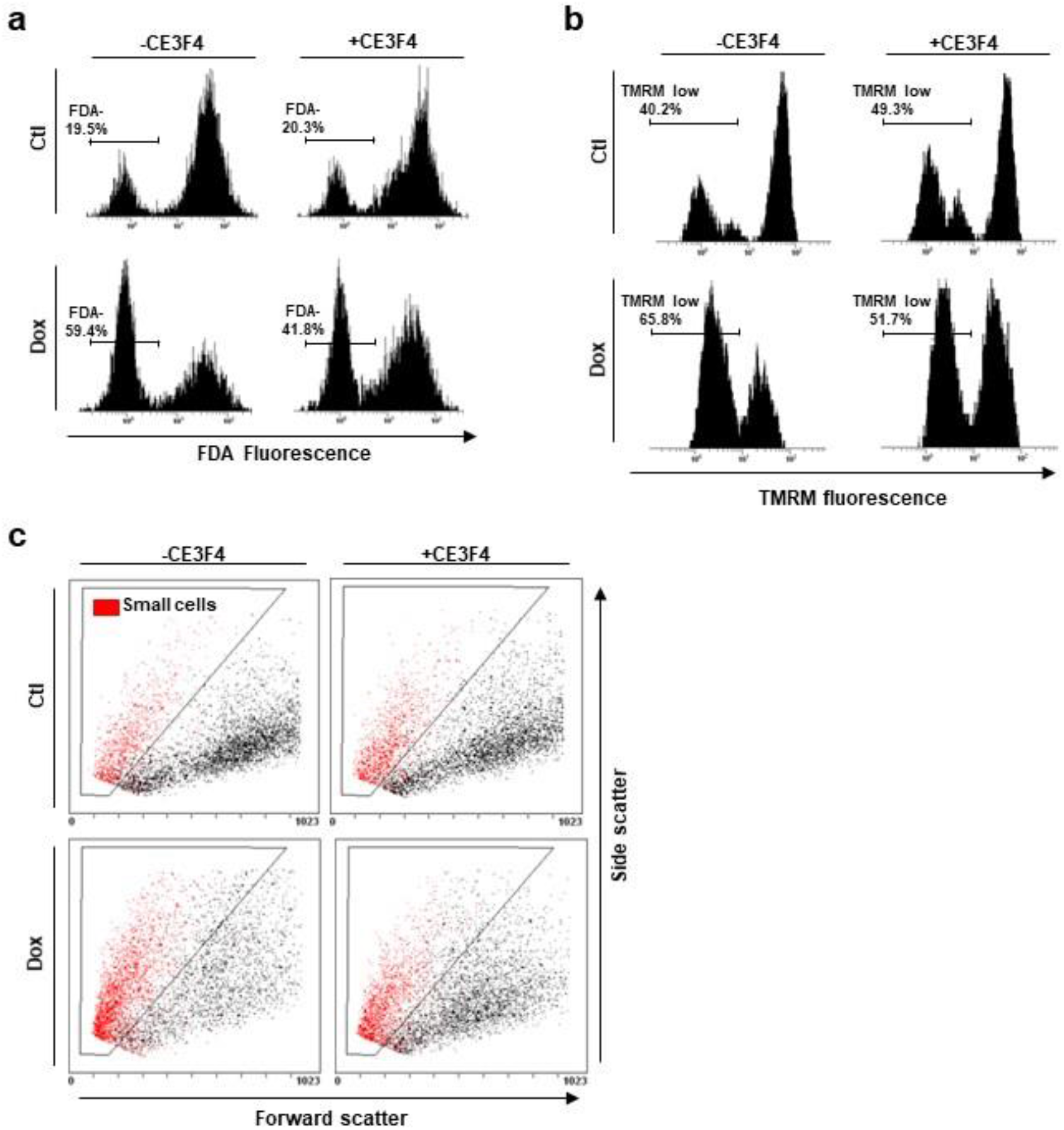
Dox induced apoptotic cell death in cardiac myocytes. **a to c,** NRVM were left untreated or treated with Dox (1 μM) +/- CE3F4 (10 μM) for 24 h and analyzed by flow cytometry. Representative monoparametric histograms of **a**, FDA fluorescence (cell death) or **b**, TMRM fluorescence (ΔΨm potential). **c**, Representative biparametric cytograms where Forward scatter signal (cell size) is represented versus Side Scatter signal (cell density).

**Extended data Fig. 2.**
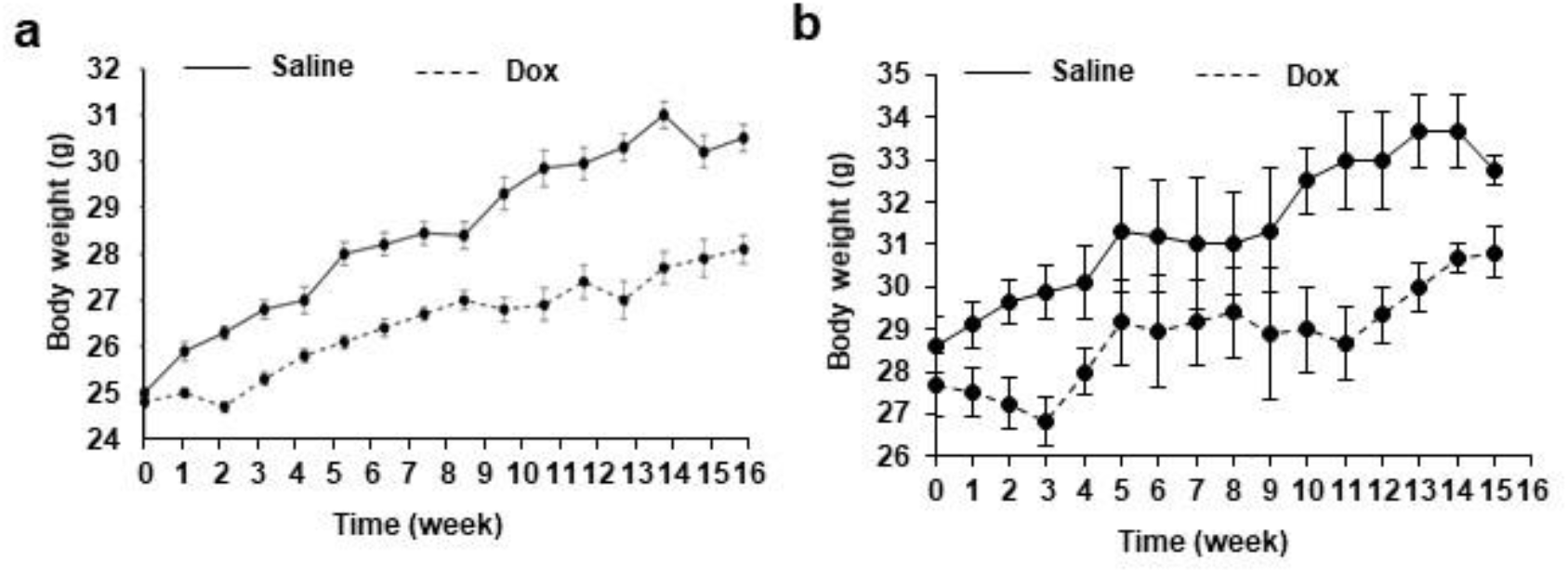
Body weight of mice treated with Dox. **a**, WT, and **b**, EPAC1 KO mice were injected (*i.v*) three times with Saline Solution (Sal) or Doxorubicin (Dox) at 4 mg/kg for each injection (10 ± 2 mg/kg cumulative dose). The body weight of mice was measured every week after treatment with Sal or Dox.

## References

1. McGowan, J.V. et al. Anthracycline Chemotherapy and Cardiotoxicity. Cardiovascular drugs and therapy 31, 63–75 (2017).

2. Lipshultz, S.E. et al. Assessment of dexrazoxane as a cardioprotectant in doxorubicin-treated children with high-risk acute lymphoblastic leukaemia: long-term follow-up of a prospective, randomised, multicentre trial. The Lancet. Oncology 11, 950–961 (2010).

3. L’Ecuyer, T. et al. DNA damage is an early event in doxorubicin-induced cardiac myocyte death. American journal of physiology. Heart and circulatory physiology 291, H1273–1280 (2006).

4. Cappetta, D. et al. Oxidative Stress and Cellular Response to Doxorubicin: A Common Factor in the Complex Milieu of Anthracycline Cardiotoxicity. Oxidative medicine and cellular longevity 2017, 1521020 (2017).

5. Rochette, L. et al. Anthracyclines/trastuzumab: new aspects of cardiotoxicity and molecular mechanisms. Trends in pharmacological sciences 36, 326–348 (2015).

6. Tokarska-Schlattner, M., Zaugg, M., Zuppinger, C., Wallimann, T. & Schlattner, U. New insights into doxorubicin-induced cardiotoxicity: the critical role of cellular energetics. Journal of molecular and cellular cardiology 41, 389–405 (2006).

7. Liu, J., Mao, W., Ding, B. & Liang, C.S. ERKs/p53 signal transduction pathway is involved in doxorubicin-induced apoptosis in H9c2 cells and cardiomyocytes. American journal of physiology. Heart and circulatory physiology 295, H1956–1965 (2008).

8. Zhang, Y.W., Shi, J., Li, Y.J. & Wei, L. Cardiomyocyte death in doxorubicin-induced cardiotoxicity. Archivum immunologiae et therapiae experimentalis 57, 435–445 (2009).

9. Casey, T.M., Arthur, P.G. & Bogoyevitch, M.A. Necrotic death without mitochondrial dysfunction-delayed death of cardiac myocytes following oxidative stress. Biochimica et biophysica acta 1773, 342–351 (2007).

10. Goormaghtigh, E., Huart, P., Praet, M., Brasseur, R. & Ruysschaert, J.M. Structure of the adriamycin-cardiolipin complex. Role in mitochondrial toxicity. Biophysical chemistry 35, 247–257 (1990).

11. Guo, J. et al. Cardioprotection against doxorubicin by metallothionein Is associated with preservation of mitochondrial biogenesis involving PGC-1alpha pathway. European journal of pharmacology 737, 117–124 (2014).

12. Kavazis, A.N., Morton, A.B., Hall, S.E. & Smuder, A.J. Effects of doxorubicin on cardiac muscle subsarcolemmal and intermyofibrillar mitochondria. Mitochondrion 34, 9–19 (2017).

13. Dorn, G.W., 2nd, Vega, R.B. & Kelly, D.P. Mitochondrial biogenesis and dynamics in the developing and diseased heart. Genes & development 29, 1981–1991 (2015).

14. Zhang, S. et al. Identification of the molecular basis of doxorubicin-induced cardiotoxicity. Nature medicine 18, 1639–1642 (2012).

15. Sawicki, K.T. et al. Preventing and Treating Anthracycline Cardiotoxicity: New Insights. Annu Rev Pharmacol Toxicol 61, 309–332 (2021).

16. Swain, S.M. et al. Cardioprotection with dexrazoxane for doxorubicin-containing therapy in advanced breast cancer. Journal of clinical oncology: official journal of the American Society of Clinical Oncology 15, 1318–1332 (1997).

17. Reichardt, P., Tabone, M.D., Mora, J., Morland, B. & Jones, R.L. Risk-benefit of dexrazoxane for preventing anthracycline-related cardiotoxicity: re-evaluating the European labeling. Future oncology (2018).

18. Tebbi, C.K. et al. Dexrazoxane-associated risk for acute myeloid leukemia/myelodysplastic syndrome and other secondary malignancies in pediatric Hodgkin’s disease. Journal of clinical oncology: official journal of the American Society of Clinical Oncology 25, 493–500 (2007).

19. Metrich, M. et al. Epac mediates beta-adrenergic receptor-induced cardiomyocyte hypertrophy. Circulation research 102, 959–965 (2008).

20. Morel, E. et al. cAMP-binding protein Epac induces cardiomyocyte hypertrophy. Circulation research 97, 1296–1304 (2005).

21. Robichaux, W.G., 3rd & Cheng, X. Intracellular cAMP sensor EPAC: Physiology, pathophysiology, and therapeutics development. Physiol Rev 98, 919–1053 (2018).

22. de Rooij, J. et al. Epac is a Rap1 guanine-nucleotide-exchange factor directly activated by cyclic AMP. Nature 396, 474–477 (1998).

23. Kawasaki, H. et al. A family of cAMP-binding proteins that directly activate Rap1. Science 282, 2275–2279 (1998).

24. Pereira, L. et al. Epac enhances excitation-transcription coupling in cardiac myocytes. Journal of molecular and cellular cardiology 52, 283–291 (2012).

25. Cazorla, O., Lucas, A., Poirier, F., Lacampagne, A. & Lezoualc’h, F. The cAMP binding protein Epac regulates cardiac myofilament function. Proceedings of the National Academy of Sciences of the United States of America 106, 14144–14149 (2009).

26. Oestreich, E.A. et al. Epac and phospholipase Cepsilon regulate Ca2+ release in the heart by activation of protein kinase Cepsilon and calcium-calmodulin kinase II. The Journal of biological chemistry 284, 1514–1522 (2009).

27. Mangmool, S., Hemplueksa, P., Parichatikanond, W. & Chattipakorn, N. Epac is required for GLP-1R-mediated inhibition of oxidative stress and apoptosis in cardiomyocytes. Molecular endocrinology 29, 583–596 (2015).

28. Okumura, S. et al. Epac1-dependent phospholamban phosphorylation mediates the cardiac response to stresses. The Journal of clinical investigation 124, 2785–2801 (2014).

29. Suzuki, S. et al. Differential roles of Epac in regulating cell death in neuronal and myocardial cells. J Biol Chem 285, 24248–24259 (2010).

30. Sag, C.M., Kohler, A.C., Anderson, M.E., Backs, J. & Maier, L.S. CaMKII-dependent SR Ca leak contributes to doxorubicin-induced impaired Ca handling in isolated cardiac myocytes. J Mol Cell Cardiol 51, 749–759 (2011).

31. Riganti, C. et al. Activation of nuclear factor-kappa B pathway by simvastatin and RhoA silencing increases doxorubicin cytotoxicity in human colon cancer HT29 cells. Mol Pharmacol 74, 476–484 (2008).

32. Huelsenbeck, S.C. et al. Rac1 protein signaling is required for DNA damage response stimulated by topoisomerase II poisons. J Biol Chem 287, 38590–38599 (2012).

33. Fazal, L. et al. Multifunctional Mitochondrial Epac1 Controls Myocardial Cell Death. Circulation research 120, 645–657 (2017).

34. Monceau, V. et al. Epac contributes to cardiac hypertrophy and amyloidosis induced by radiotherapy but not fibrosis. Radiotherapy and oncology: journal of the European Society for Therapeutic Radiology and Oncology 111, 63–71 (2014).

35. Huk, D.J., Ashtekar, A., Magner, A., La Perle, K. & Kirschner, L.S. Deletion of Rap1b, but not Rap1a or Epac1, Reduces Protein Kinase A-Mediated Thyroid Cancer. Thyroid: official journal of the American Thyroid Association 28, 1153–1161 (2018).

36. Kumar, N. et al. Insights into exchange factor directly activated by cAMP (EPAC) as potential target for cancer treatment. Molecular and cellular biochemistry (2018).

37. Jansen, S.R. et al. Epac1 links prostaglandin E2 to beta-catenin-dependent transcription during epithelial-to-mesenchymal transition. Oncotarget 7, 46354–46370 (2016).

38. Almahariq, M. et al. Pharmacological inhibition and genetic knockdown of exchange protein directly activated by cAMP 1 reduce pancreatic cancer metastasis in vivo. Molecular pharmacology 87, 142–149 (2015).

39. Almahariq, M., Mei, F.C. & Cheng, X. The pleiotropic role of exchange protein directly activated by cAMP 1 (EPAC1) in cancer: implications for therapeutic intervention. Acta biochimica et biophysica Sinica 48, 75–81 (2016).

40. Wang, P. et al. Exchange proteins directly activated by cAMP (EPACs): Emerging therapeutic targets. Bioorganic & medicinal chemistry letters 27, 1633–1639 (2017).

41. Courilleau, D. et al. Identification of a tetrahydroquinoline analog as a pharmacological inhibitor of the cAMP-binding protein Epac. The Journal of biological chemistry 287, 44192–44202 (2012).

42. Zhu, Y. et al. Biochemical and pharmacological characterizations of ESI-09 based EPAC inhibitors: defining the ESI-09 “therapeutic window”. Scientific reports 5, 9344 (2015).

43. Tsalkova, T. et al. Isoform-specific antagonists of exchange proteins directly activated by cAMP. Proceedings of the National Academy of Sciences of the United States of America 109, 18613–18618 (2012).

44. Moro, S. et al. Interaction model for anthracycline activity against DNA topoisomerase II. Biochemistry 43, 7503–7513 (2004).

45. Llach, A. et al. Progression of excitation-contraction coupling defects in doxorubicin cardiotoxicity. J Mol Cell Cardiol 126, 129–139 (2019).

46. Li, M. et al. Phosphoinositide 3-Kinase Gamma Inhibition Protects from Anthracycline Cardiotoxicity and Reduces Tumor Growth. Circulation (2018).

47. Zhang, Y. et al. A novel agent attenuates cardiotoxicity and improves antitumor activity of doxorubicin in breast cancer cells. Journal of cellular biochemistry (2018).

48. Lim, C.C. et al. Anthracyclines induce calpain-dependent titin proteolysis and necrosis in cardiomyocytes. The Journal of biological chemistry 279, 8290–8299 (2004).

49. Lebrecht, D. & Walker, U.A. Role of mtDNA lesions in anthracycline cardiotoxicity. Cardiovascular toxicology 7, 108–113 (2007).

50. Singhmar, P. et al. Orally active Epac inhibitor reverses mechanical allodynia and loss of intraepidermal nerve fibers in a mouse model of chemotherapy-induced peripheral neuropathy. Pain 159, 884–893 (2018).

51. Childs, A.C., Phaneuf, S.L., Dirks, A.J., Phillips, T. & Leeuwenburgh, C. Doxorubicin treatment in vivo causes cytochrome C release and cardiomyocyte apoptosis, as well as increased mitochondrial efficiency, superoxide dismutase activity, and Bcl-2:Bax ratio. Cancer research 62, 4592–4598 (2002).

52. Deniaud, A. et al. Endoplasmic reticulum stress induces calcium-dependent permeability transition, mitochondrial outer membrane permeabilization and apoptosis. Oncogene 27, 285–299 (2008).

53. Carvalho, F.S. et al. Doxorubicin-induced cardiotoxicity: from bioenergetic failure and cell death to cardiomyopathy. Medicinal research reviews 34, 106–135 (2014).

54. Wang, H. et al. Inhibition of Epac1 suppresses mitochondrial fission and reduces neointima formation induced by vascular injury. Sci Rep 6, 36552 (2016).

55. Corbett, K.D. & Berger, J.M. Structure, molecular mechanisms, and evolutionary relationships in DNA topoisomerases. Annual review of biophysics and biomolecular structure 33, 95–118 (2004).

56. Nitiss, J.L. Targeting DNA topoisomerase II in cancer chemotherapy. Nature reviews. Cancer 9, 338–350 (2009).

57. Baljinnyam, E. et al. Epac1 increases migration of endothelial cells and melanoma cells via FGF2-mediated paracrine signaling. Pigment cell & melanoma research 27, 611–620 (2014).

58. Baljinnyam, E. et al. Epac1 promotes melanoma metastasis via modification of heparan sulfate. Pigment cell & melanoma research 24, 680–687 (2011).

59. Gao, M. et al. Epac1 knockdown inhibits the proliferation of ovarian cancer cells by inactivating AKT/Cyclin D1/CDK4 pathway in vitro and in vivo. Medical oncology 33, 73 (2016).

60. Kumar, N., Gupta, S., Dabral, S., Singh, S. & Sehrawat, S. Role of exchange protein directly activated by cAMP (EPAC1) in breast cancer cell migration and apoptosis. Molecular and cellular biochemistry 430, 115–125 (2017).

61. Laurent, A.C. et al. Exchange protein directly activated by cAMP 1 promotes autophagy during cardiomyocyte hypertrophy. Cardiovasc Res 105, 55–64 (2015).

62. Desai, V.G. et al. Development of doxorubicin-induced chronic cardiotoxicity in the B6C3F1 mouse model. Toxicology and applied pharmacology 266, 109–121 (2013).

63. Rochais, F. et al. Negative feedback exerted by cAMP-dependent protein kinase and cAMP phosphodiesterase on subsarcolemmal cAMP signals in intact cardiac myocytes: an in vivo study using adenovirus-mediated expression of CNG channels. The Journal of biological chemistry 279, 52095–52105 (2004).

64. Courilleau, D., Bouyssou, P., Fischmeister, R., Lezoualc’h, F. & Blondeau, J.P. The (R)-enantiomer of CE3F4 is a preferential inhibitor of human exchange protein directly activated by cyclic AMP isoform 1 (Epac1). Biochemical and biophysical research communications 440, 443–448 (2013).

65. Maillet, M. et al. Crosstalk between Rap1 and Rac regulates secretion of sAPPalpha. Nature cell biology 5, 633–639 (2003).

66. Brown, L.M., Rogers, K.E., McCammon, J.A. & Insel, P.A. Identification and validation of modulators of exchange protein activated by cAMP (Epac) activity: structure-function implications for Epac activation and inhibition. The Journal of biological chemistry 289, 8217–8230 (2014).

67. Wartlick, F., Bopp, A., Henninger, C. & Fritz, G. DNA damage response (DDR) induced by topoisomerase II poisons requires nuclear function of the small GTPase Rac. Biochimica et biophysica acta 1833, 3093–3103 (2013).

